# Unsupervised anomaly detection for tumor delineation in a preclinical model of glioblastoma using CEST MRI

**DOI:** 10.64898/2026.02.17.706435

**Authors:** Anshuman Swain, Abeer Mathur, Narayan Datt Soni, Neil Wilson, Blake Benyard, Paul Jacobs, Sunil Kumar Khokhar, Dushyant Kumar, Mohammad Haris, Ravinder Reddy

**Author notes:** Corresponding author:* Ravinder Reddy. Anshuman Swain and Abeer Mathur contributed equally to this work.

## Abstract

**Introduction:** Glioblastoma is characterized by heterogeneous tumor characteristics and infiltrative tumor boundaries, making accurate delineation difficult with extensive manual annotations. Chemical exchange saturation transfer (CEST) is a non-invasive MRI technique used for *in* vivo assessment of metabolic and macromolecular information through a Z-spectrum. CEST may provide insight into metabolic changes present in early-stage disease that are not visible in routine clinical imaging, thereby improving tumor delineation. In this work, we use an unsupervised anomaly detection (UAD) strategy to learn the distribution of features present in Z-spectra of healthy tissue and capture their deviations in pathology, foregoing the need for extensive labels. The approach leverages the metabolic information provided by CEST to improve the detection and delineation of glioblastoma and inform further treatment planning.

**Methods:** A 1D convolutional autoencoder (CAE) was implemented to reconstruct Z-spectra from individual tissue voxels. The network was trained on Z-spectra acquired at 9.4T from healthy Sprague-Dawley rats and tested on data acquired from F98 glioma-bearing rats post Gd-administration. For baseline comparisons, Isolation Forest and Local Outlier Factor, which have shown success in anomaly detection, were implemented. For the CAE, our anomaly score was determined to be the mean squared reconstruction error. To facilitate clinical translation and evaluate the robustness of our model for under sampled Z-spectra, acceleration factors of 2x and 7x were performed with two sampling schemes: uniformly skipping frequency offsets and selecting offsets based on feature importance identified by Shapley value analysis and Integrated Gradients (IG). Binarization was performed by determining an optimal anomaly threshold, followed by comparison to ground truth tumor masks. Metrics related to model performance were assessed for baseline anomaly detectors on the fully sampled dataset and for the CAE on fully and under sampled datasets.

**Results:** The best baseline anomaly detector was Isolation Forest, with an ROC-AUC of 0.967 and an F1-score of 0.584. Our method, the CAE, accurately reconstructed Z-spectral features, achieving Dice scores of up to 0.72 and outperforming the baseline model with an ROC-AUC of 0.968 and F1-score of 0.642. This model performance remained robust across sampling schemes and acceleration factors, with ROC-AUCs of ∼0.96 and similar Dice scores (up to 0.7). Feature importance analysis indicated that offsets in the range of ±3.0 to 5.0ppm contributed most to the anomaly score.

**Discussion:** This study successfully demonstrated a UAD pipeline utilizing the Z-spectrum from CEST MRI for metabolically informed tumor delineation. The framework captures biochemical deviations that may precede or extend beyond morphologic abnormalities, enabling sensitive detection of tumor regions and intra-tumoral heterogeneity that previous methods may fail to capture. The offsets from the feature analysis indicated a strong contribution from the magnetization transfer (MT) pool to the spectral deviations captured by the model, with additional contributions from relayed nuclear Overhauser effect (rNOE) and amide proton transfer (APT). Model robustness with under sampling further highlights the pipeline’s potential in accelerated acquisitions, thus improving clinical practicality. While there is a need for validation on larger cohorts and clinical datasets, the current results demonstrate that this label-free, Z-spectral anomaly mapping can serve as an interpretable and scalable tool for monitoring tumor heterogeneity and progression, with potential applicability to other diffuse or metabolically subtle pathologies.

## Introduction

Chemical Exchange Saturation Transfer (CEST) is an MR imaging method that can non-invasively detect metabolites and macromolecules *in vivo* with high sensitivity. CEST exploits the exchange between solute protons and bulk water protons to provide indirect measures for a range of endogenous and exogenous metabolites (1, 2). Analysis of the Z-spectrum acquired in CEST is often conducted by performing multi-pool Lorentzian fitting (MPLF), providing semi-quantitative estimates of metabolite concentrations (3, 4). Metrics derived from Z-spectral analysis, such as amide proton transfer (APT) and relayed nuclear Overhauser effect (rNOE), have been employed in both clinical and preclinical studies, revealing metabolic underpinnings and dysfunction in disease and pathological tissue types, like lesions and tumors (5, 6). CEST can also be used to assess microenvironment properties such as pH, temperature, and oxidative stress (7). Despite the utility of multi-pool fitting, the technique is ill-posed with solutions varying based on parameter initializations and constraints (8). When performing comparisons between healthy and pathological conditions, sufficiently large sample sizes are required, with analyses performed in a group-wise manner (e.g., healthy subjects v. Alzheimer’s disease patients) and in pre-defined ROIs, which can obscure tissue heterogeneity. In addition, defining distinct sub-groups proves difficult in heterogeneous diseases, where individuals can present with different phenotypic manifestations at similar stages of pathology. A prior clinical diagnosis is also necessary for defining populations, making it challenging to group individuals in early stages of the disease. Consequently, detecting early-stage disease pathology and monitoring its progression on an individual basis is vital.

Existing unsupervised anomaly detection (UAD) strategies have demonstrated increased potential for detecting and monitoring disease progression on an individual basis (9, 10). Machine learning pipelines, including deep learning approaches, have been developed to identify anomalies in the context of images, tabular data, and time series (11). The strength of UAD stems from its ability to identify anomalies without access to extensive labeled data (12). Consequently, only healthy datasets, which are much easier to obtain and more readily accessible, are required to train the model. When tested on unseen data, samples that significantly deviate from the training distribution (i.e., healthy/normal data) are classified as anomalies. UAD has shown success in medical image anomaly detection, where models are trained to reconstruct healthy image data (13). Upon testing with images that contain pathological tissue, the model generates a ‘healthy’ version of this image and metrics like mean squared or mean absolute error used to identify anomalies.

Our goal is to combine UAD with CEST to identify anomalies in a rat glioblastoma model, which are highly heterogeneous (14). Our methodology operates on the Z-spectrum rather than on an image, thereby deviating from previous medical image anomaly detection approaches by attempting to identify anomalous metabolic signatures instead of largely morphological changes observed through conventional imaging techniques. In addition, since each voxel in an image has a corresponding Z-spectrum, we can greatly amplify our training size with a minimal number of subjects. For example, a 3D acquisition performed on one subject can provide over 100k training samples. In addition, the sensitivity of CEST to early-stage metabolic changes may enable us to potentially detect anomalies prior to the appearance of morphological abnormalities (15, 16), conferring advantages over available methodologies that rely primarily on structural and anatomic images (17). We extend previous work from our group, which implemented UAD (18) on an MS patient, to preclinical tumor models and show the robustness of our methodology to under sampled acquisitions of Z-spectrum frequencies, thus improving clinical feasibility.

## Methods

### 1.1. Tumor inoculation

The experimental protocols used in this study were approved by the Institutional Animal Care and Use Committee of the University of Pennsylvania. In this study, data were acquired on on the brains of four healthy and three tumor-bearing Sprague-Dawley rats (∼6 weeks old) (19). For tumor inoculation, rat heads were fixed on a stereotaxic apparatus with a continuous supply of isoflurane to induce a surgical plane of anesthesia. Bupivacaine (under the scalp; 2mg/kg) and meloxicam (subcutaneous; 2mg/kg) were provided as analgesics. The head was shaved using a trimmer, and an incision was made to expose the skull. After sterilizing the scalp, a burr hole was drilled 3mm lateral (right) and 3mm posterior to the bregma using stereotactic navigation, and a 5uL suspension containing ∼50k F98 (gliosarcoma) cells in phosphate-buffered saline (PBS) was injected into the cerebral cortex (2mm deep) at a rate of 0.5 μL/min using a 10 μL Hamilton syringe mounted with 32-gauge needle and an infusion pump (Stoelting Co, USA). Sutures were used to close the scalp’s incision, and the rats were monitored every day to assess wound healing and surgical complications.

### 1.2. Data Acquisition

To monitor the size of the tumor, rats (n = 3) were scanned every week using T_2_-weighted rapid-acquisition rapid-echo (rapid-acquisition rapid-echo; 16 slices, TE1/TE2/TE3 = 15/45/75 ms, TR = 3s, and two averages) and T1-weighted FLASH (fast low-angle shot; 16 slices, TE=4 ms, TR=200 ms, and four averages) images. Once the tumor was clearly visible with minimal indications of necrosis, contrast-enhanced and CEST acquisitions were performed as follows:

First, a localizer was acquired to position the brain, followed by T_1_-weighted FLASH and T_2_-weighted RARE images. For contrast-weighted images, a Gadolinium-based contrast agent (Gd-DOTA, Gadoerate meglumin, Dotarem, France; 0.1 mmol/kg) was administered intravenously through a catheter tail vein and T_1_-weighted images were acquired every 2 minutes. Once the signal enhancement reached its peak (∼10 minutes), the CEST acquisition was performed on a slice of interest. To correct for B0 inhomogeneities, a WASSR (WAter Saturation Shift Referencing (20); TE = 4 ms, TR 410 ms, 22 frequency offsets from 0 to ±1 ppm in steps of 0.1ppm, B1rms = 0.1μT) was acquired. A full Z-spectrum was acquired for the CEST-weighted images with 52 offsets ranging from +5 to -5 ppm in steps of 0.2 ppm. The acquisition parameters were TE= 4ms, TR = 3s, B1 = 1.0 μT, saturation duration (tsat) = 3s, and two averages. An unsaturated (i.e., reference) image (with the same parameters as the saturated images, but with B1 = 0.0uT and an offset of -300ppm) was additionally acquired. For all images, the FOV was 30 mm x 30 mm with an image matrix size of 192 × 192, resulting in an in-plane resolution of 0.156 mm x 0.156 mm for all images. For healthy rats (n = 4), Gd-administration was not performed and T_1_ and T_2_-weighted images were acquired only once for slice selection, followed by the CEST acquisition.

### 1.3. Image post-processing, metabolite quantification, and data preparation

For healthy rats, the slice of interest from the T_2_-weighted image was skull-stripped using a fuzzy c-means clustering algorithm, followed by a series of erosions and dilations on the brain mask to obtain the final brain segmentation. For tumor-bearing rats, the brain was manually segmented using ITK-SNAP (21) to avoid issues with the tumor region. To generate the ground-truth tumor mask, three lab members were asked to perform the tumor segmentation on the Gd-enhanced T_1_-weighted and T_2_-weighted images, and a majority vote was used to determine the final tumor mask. The selected lab members (A.S., A.M., and N.D.S.) have expertise in visualizing mouse brain anatomy, and segmentations between the members were largely similar in the tumor core with discrepancies at tumor boundaries. Z-spectrum images were generated by normalizing the CEST-weighted images by the reference image. Denoising was performed on normalized Z-spectra by exploiting redundancies in saturation offsets using singular value decomposition (SVD). Briefly, the image matrix was reshaped to [voxels x offsets], followed by SVD, selection of the singular value threshold based on the median criterion, and recovery of the low-rank, denoised data (22, 23). Voxel-wise multi-pool Lorentzian fitting was performed with five metabolite pools: direct saturation (DS), magnetization transfer (MT), amide proton transfer (APT), amine, and relayed Nuclear Overhauser effect (NOE) using initial parameter estimates and bounds from Windschuch et. al. (24). For subsequent anomaly detection, two datasets were constructed for healthy and tumor rats: (1) (1 – Z-spectrum) used as input to the CAE model and classical ML anomaly detection models, and (2) the fitted parameters, including chemical shift, amplitude, and linewidth, derived from multi-pool fitting as input to classical ML anomaly detection models. For the first dataset, the image matrix was reshaped from [height x width x offsets] to [voxels x offsets], while the second dataset was of shape [voxels x estimated parameters]. Both datasets were designed to evaluate anomaly detection performance when feature information is represented as offsets versus fitted parameters. The data from healthy rats was concatenated to get the final training set. For tumor-bearing rats, one rat was held out as a validation set to determine the anomaly threshold, while the other two rats were used for testing.

### 1.4 Model selection and implementation

#### 1.4.1. Local outlier factor (LOF) and Isolation Forest (IF)

Two datasets, as stated prior, were used for baseline machine learning methods: one dataset treated the fitted parameters from the Lorentzian-analysis of the Z-spectrum as features, while the other dataset used the Z-spectrum frequency offsets as features. Local outlier factor (LOF) and isolation forest (IF), implemented in Scikit-Learn (25), served as baseline anomaly detectors. LOF (26) compares the density of a point to the density of its nearest neighbors, with outliers (i.e., anomalies) expected to exist in a sparse region of the feature space while inliers exist in denser areas. IF (27) builds an ensemble of decision trees, and samples that require fewer splits to be separated are more likely to be anomalies. A hyperparameter search was performed for each model by selecting the hyperparameters that maximized the ROC-AUC and F1-score on the validation set. The optimal anomaly threshold for each model, derived from the highest F1-score, was used for subsequent evaluations on the test set. Anomaly score maps were generated for comparison to our methodology and ground truth labels.

#### 1.4.2. Convolutional autoencoder (CAE)

Our proposed model in this study was a convolutional autoencoder. CAEs have shown success in several anomaly detection tasks due to their local inductive bias and ability to capture neighborhood correlations (28). Their use and success in time series anomaly detection provided the foundation for our implementation (29). For our task, we implemented a 1D CAE, in which the encoder captures local spectral information and generates an encoded representation (i.e., latent representation) of this information. The decoder then uses the representation to reconstruct the spectrum (**Figure 1**). Our model was trained on an NVIDIA RTX 4090 graphics processing unit for 150 epochs with an early stopping criterion (∼40 epochs) using the Stochastic Gradient Descent optimizer and a learning rate of 1e-4. The anomaly score is calculated as the mean squared error between the reconstructed and input spectra. Higher reconstruction errors are anticipated in pathological voxels since the CAE only learns the distribution and manifold of healthy data. Voxel-wise anomaly scores are reshaped back into the image space, and the threshold for binarization of the test set is set as the 90^th^ percentile of anomaly scores in the validation set (i.e., one of the tumor-bearing rats). The threshold was empirically determined by maximizing the Dice score on the validation set. Although the threshold selection is performed in a supervised manner, the training paradigm is entirely unsupervised, and labels are not used for model optimization. The proposed approach follows a standard unsupervised anomaly detection strategy with supervised threshold calibration (30, 31).

**Figure 1.**
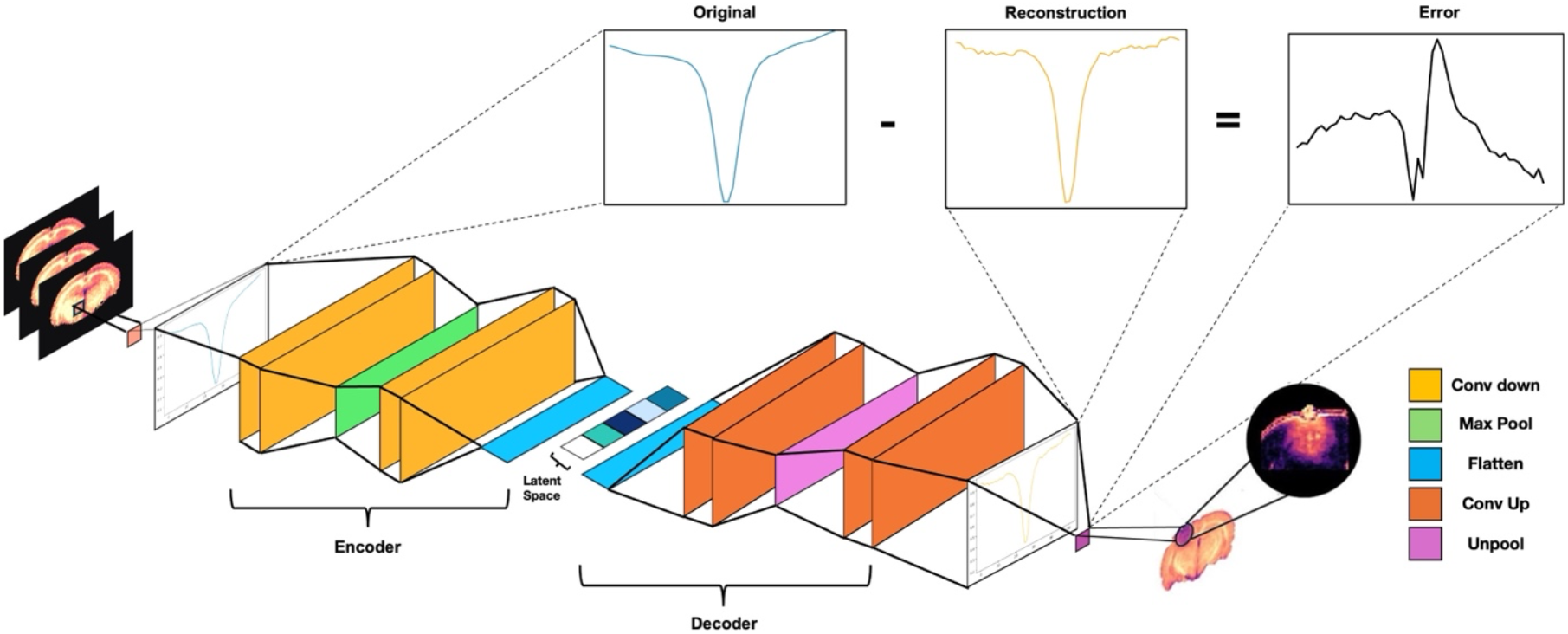
Schematic representation of a 1D Convolutional Autoencoder. Each voxel, bearing a full Z-spectrum, is fed into an encoder which generates a latent representation that captures the most important features from the (1 - Z-spectrum). A symmetric decoder reconstructs the spectrum from the latent space, and the mean square error (MSE) between the input and output yields the per-voxel anomaly score. The block colors denote layer types.

#### 1.4.3. Feature importance

Given that the Z-spectrum contains metabolic information, it is important to understand which offsets (i.e., features) contribute the most in identifying anomalies. To determine feature importance, Shapley values were computed for IF. In game theory, the Shapley value is a method of fairly distributing the total payout (i.e., gains/costs) to players who collaborated to achieve a specific goal. Shapley values have been more recently adopted in ML, in which the importance of a feature in achieving the model’s goal, such as prediction, is analogous to a player’s contribution in winning a cooperative game (32). To implement Shapley values, a baseline, which describes the average outcome of a specific model, is needed. For the IF, the baseline was defined as the distribution of anomaly scores of healthy voxels. Shapley values would thus indicate whether an offset increases or decreases deviation from the distribution of healthy tissue. If a feature contributed to a higher anomaly score, a more negative Shapley value was expected since negative values from the IF’s decision function indicate more anomalous instances.

In addition to Shapley values, we implemented integrated gradients (IG) (33) for the CAE as this method is well-suited for differentiable, deep learning models and can capture the complex, non-linear interactions that Shapley values may miss. For IG calculation, we wrapped the CAE in a function that calculated the MSE between the input and output as the models’ final prediction. IG determines feature importance in our CAE by integrating the gradient of the MSE, with respect to each offset, from the baseline to the input Z-spectrum. A larger integrated gradient for an offset indicates a greater sensitivity of the MSE (i.e., anomaly score) to changes in that particular offset. Following the calculation of IG for our CAE, we normalized the path values for each offset by the mean Z-spectrum value for that particular offset. This normalization is performed since the gradients are sensitive to the absolute signal of an offset in the Z-spectrum as well as the gradient magnitude, as given by the following equation:

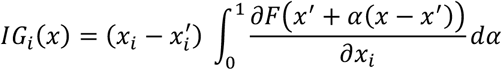

Consequently, offsets with higher amplitudes and regions where sharp transitions in intensity occur, especially near direct saturation at 0ppm, may exhibit higher IG values that are not driven by changes in tumor pathology. We selected the sub-sampled (i.e., feature importance guided) offsets using the normalized IG values. Analysis of the average Z-spectrum from healthy and tumor voxels revealed MT-driven amplitude shifts as the dominant feature, so priority was given to those offsets that captured MT pool contributions.

#### 1.4.5. Metrics for model evaluation

The following metrics were used to evaluate baseline anomaly detectors and the CAE: accuracy, F1-score, ROC-AUC, PR-AUC, precision, and recall. In addition, for the CAE, Dice scores were calculated to evaluate the degree of overlap between the binarized model segmentations and the ground truth segmentations for each rat in the test set (i.e., Dice 1 and Dice 2).

#### 1.4.4. Sub-sampling of Z-spectrum

Since the acquisition of a Z-spectrum is time consuming and limits its clinical utility, we trained the CAE on sub-sampled sets of frequency offsets and calculated the aforementioned metrics. We utilized two sub-sampling schemes: the first scheme uniformly selected offsets based on the acceleration factor (i.e., 2x acceleration resulting in selection of every other offset), while the second scheme selected the top ‘k’ features based on their importance, as determined by IG. The two acceleration factors that were evaluated were 2x and 7x.

## Results

### 2.1. LOF and IF

Metrics from LOF and IF are presented in Table 1.

**Table 1.** Metrics from the CAE and baseline anomaly detection models—IF and LOF—when using the Z-spectrum offsets or fitted parameters from multi-pool Lorentzian fitting as features. The CAE exhibits the best performance metrics except for Recall, in which the IF trained on the Z-spectrum exhibits moderately higher performance.

| Model | Accuracy | F1 | ROC AUC | PR AUC | Precision | Recall |
| --- | --- | --- | --- | --- | --- | --- |
| **CAE** | **0.929** | **0.642** | **0.968** | **0.522** | **0.493** | 0.921 |
| **IF: Z-spectrum** | 0.902 | 0.584 | 0.967 | 0.512 | 0.414 | 0.994 |
| **LOF: Z-spectrum** | 0.418 | 0.192 | 0.749 | 0.125 | 0.106 | **0.997** |
| **IF: Fitted parameters** | 0.692 | 0.256 | 0.776 | 0.142 | 0.154 | 0.761 |
| **LOF: Fitted parameters** | 0.775 | 0.289 | 0.797 | 0.182 | 0.185 | 0.656 |

Overall, LOF and IF performed poorly in detecting anomalous voxels when fitted parameters were used as features, with ROC-AUCs of 0.797 and 0.776 and PR-AUCs of 0.182 and 0.142, respectively. The performance is evident in the anomaly score maps, where Rat #1 shows only mild anomaly scores (i.e., higher values) in the tumor region for both LOF and IF, while Rat #2 shows clearer anomaly features with IF compared to LOF (**Figure 2a**). In comparison, when using the Z-spectrum offsets as features, IF shows stark differences between the tumor and healthy-appearing tissue for both rats (ROC-AUC: 0.967 and PR-AUC: 0.512), while LOF fails to highlight those differences in Rat #1 and achieves a moderate detection in Rat #2 (**Figure 2b**), with ROC-AUC and PR-AUC of 0.749 and 0.125). Based on these observed differences and quantitative metrics, IF with Z-spectrum offsets as features was selected as the baseline model for subsequent comparisons.

**Figure 2.**
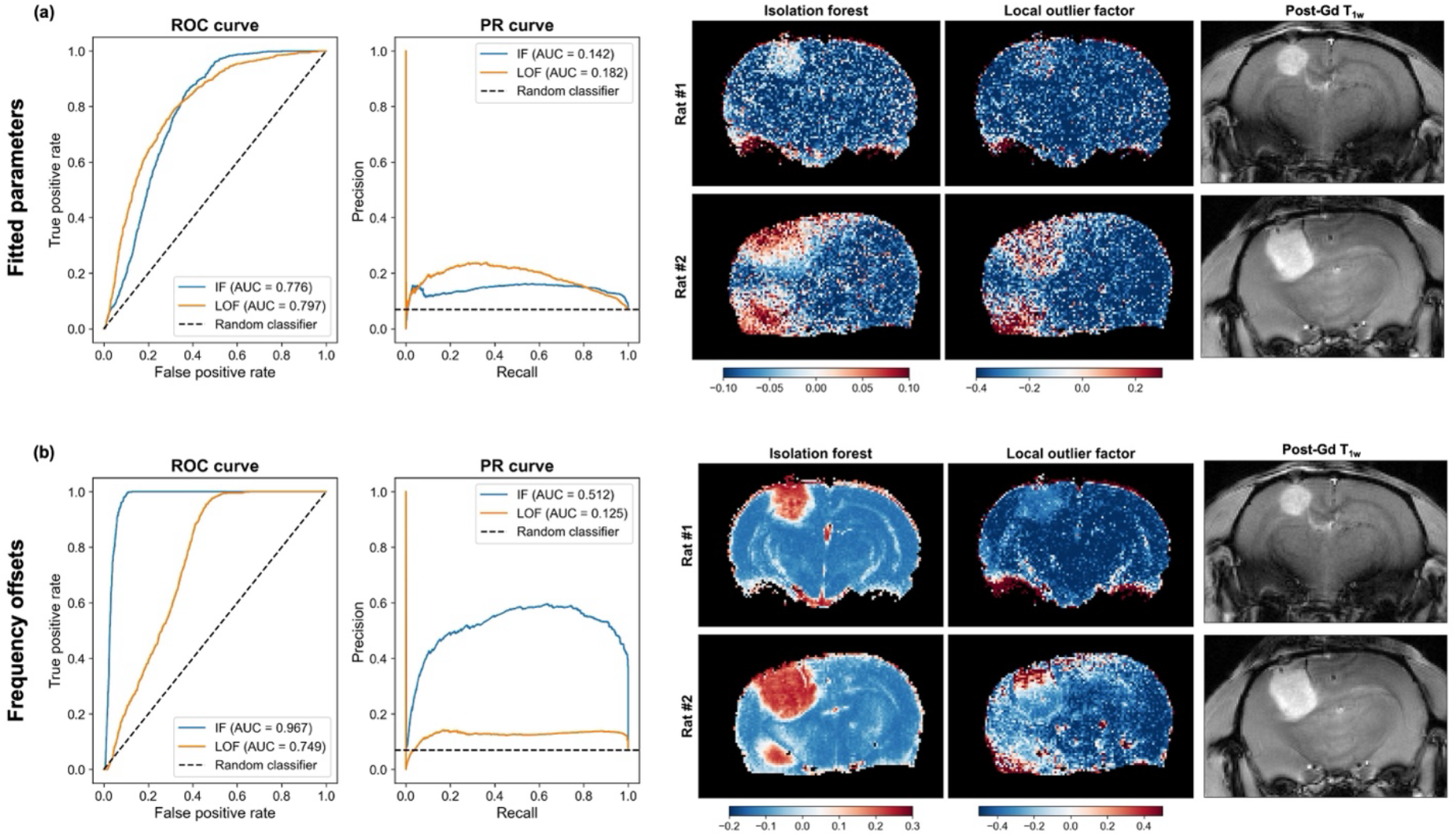
The figure above compares anomaly detection results for Isolation Forest (IF) and Local Outlier Factor (LOF) using fitted parameters or frequency offsets as features. (a) showcases the ROC and PR curve alongside anomaly score maps for the models, suggesting that the fitted parameters perform poorly as features for anomaly detection. (b) highlights the significantly improved performance of IF when using Z-spectral offsets as features, while LOF still struggles at capturing anomalous voxels. Post-contrast T_1_-weighted images are provided for anatomical reference.

### 2.2. CAE

Reconstruction residuals for tumor voxels remained systematically higher than for healthy voxels (**Figure 3a**). This pattern supports our central assumption that the latent space of a CAE trained exclusively on healthy Z-spectra defines a manifold from which tumor voxels deviate. Interestingly, tumor voxels exhibit a high reconstruction error across all offsets, with lower error observed at ∼0 ppm. This observation is consistent with results from multi-pool Lorentzian fitting (**Supplementary Figure 1(a-b)**). Differences in DS, with a chemical shift centered at 0 ppm, are less apparent between the tumor and normal-appearing tissue given the low F1-score despite a high ROC-AUC, while MT, a broad component that contributes to all offsets, shows a marked difference between these tissue types with both high ROC-AUC and F1-scores. The anomaly score maps show variation and higher error in the tumor core compared to the boundary, demonstrating the model’s ability to capture disease heterogeneity and progression (**Figure 3b**). After thresholding the anomaly scores, binary maps suitable for segmentation of the tumor region were generated and compared to the T1 weighted images and ground truth masks. The binarized masks overlapped significantly with the ground truth, yielding Dice scores of 0.5 and 0.72. However, detection of anomalous voxels around the edges of the brain – likely from dura matter – reduces the overlap, and so the reported Dice scores may be underestimated. Nonetheless, the model demonstrated robust performance in delineating tumor heterogeneity and segmenting the anomalous tumor region in each rat.

**Figure 3.**
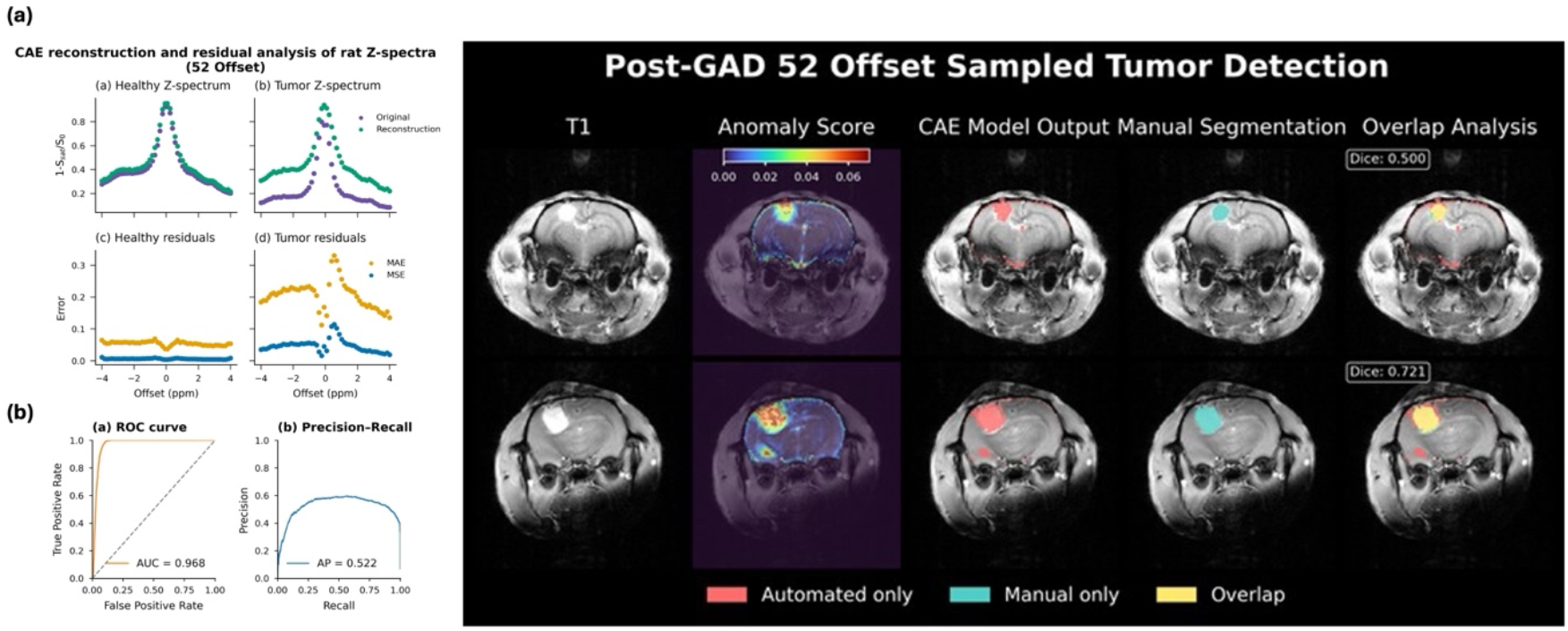
(a) Mean Z-spectrum and mean convolutional autoencoder (CAE) reconstruction are displayed for healthy and tumor voxels. Reconstruction errors between the original spectrum and reconstruction are consistently higher for tumor voxels than for healthy rats, contributing to high anomaly scores as displayed in the anomaly maps. (b) Metrics from the ROC and PR curve demonstrate the robust performance of the CAE in anomaly detection. Dice scores indicate high overlap between predicted binary tumor masks and ground truth tumor masks.

### 2.3. Feature importance and sub-sampling of Z-spectrum

The most important offsets determined by Shapley values from the IF were -3.4 and -1.2 ppm, corresponding to the chemical shifts of rNOE. Overall, the feature importance was heterogeneous, capturing changes around 0 ppm and 2 ppm, as well as offsets in the range of -3 to -5 ppm (**Figure 4a**). Normalized integrated gradient values from the CAE also highlighted offsets in the range of ±3 to ±5 ppm, in addition to highlighting offsets around 2ppm (**Figure 4b**). The offsets in these ranges typically guide the Lorentzian fit of the MT pool, which we observed to be distinctively lower in tumor tissue, as well as rNOE and amine, and so the final selection of the top ‘K’ optimal offsets was driven by a combination of model predictions and domain knowledge.

**Figure 4.**
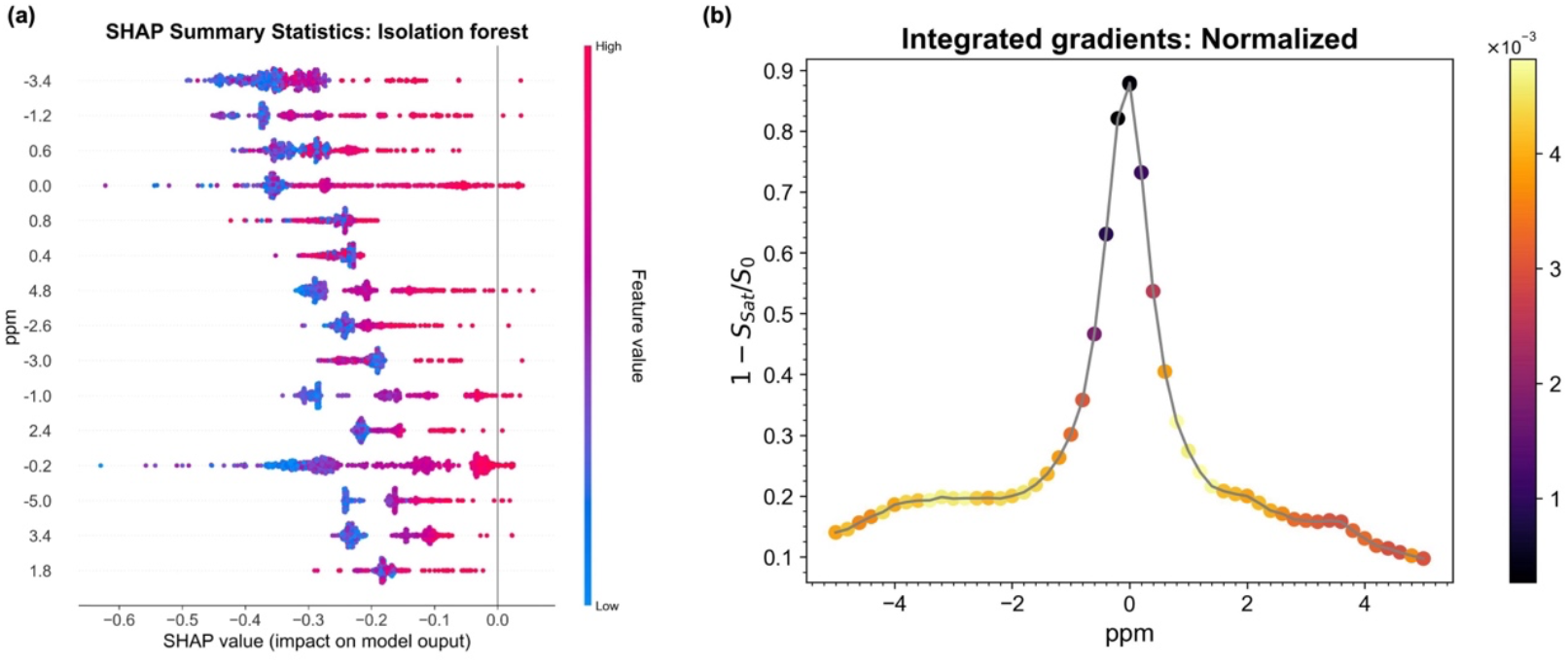
(a) Plot showing the top 15 most important features calculated by Kernel SHAP for the IF. For IF, larger negative SHAP values indicate a feature that contributes to a higher anomaly score. The two most important offsets were -3.4 and -1.2 ppm, corresponding to rNOE pool contributuions (b) Plot showing the normalized integrated gradient values derived from the CAE. Offsets in the range of -3 to -5 ppm show high IG scores, with contributions likely stemming from rNOE and MT, followed by offsets around 2 ppm which correspond to the amine pool. For selection of sub-sampled offsets, priority was placed on MT-related offsets as determined by IG feature attributions and observed changes in Z-spectrum amplitudes between tumor and healthy tissue.

Metrics from uniform and feature importance guided sub-sampling schemes are presented in **Table 2**. Across all sub-sampling schemes, the ROC-AUCs and Dice scores are similar to the fully sampled data, indicating the CAE’s ability to capture the most informative spectral information in the latent space even when features are sparse. Interestingly, the uniform sub-sampling scheme with 2x acceleration showed metrics closest to the fully sampled data. However, the feature importance sub sampling performed better at the higher acceleration factor. This is reflected in the anomaly score maps (**Figure 5**), with the uniform 2x under sampling having an almost indiscernible anomaly map compared to the fully sampled one.

**Table 2.** Metrics from the CAE trained on two under-sampling schemes with two different acceleration factors. The original number of offsets shows the best performance across all metrics as expected, albeit with slightly lower precision. Across all under-sampling schemes and factors, the 26 offset uniform sub-sampling performs the best, with the 8 offset uniform sub-sampling having slightly higher Recall and Dice 1 scores. Dice 1 and Dice 2 refer to the Dice scores of the test rats presented in the top row and bottom row of Figure 3, respectively.

| Sampling Scheme | Accuracy | F1 | ROC-AUC | PR-AUC | Precision | Recall | Dice 1 | Dice2 |
| --- | --- | --- | --- | --- | --- | --- | --- | --- |
| 52 Offsets: Original | 0.929 | 0.642 | 0.968 | 0.522 | 0.493 | 0.921 | 0.5 | 0.721 |
| **26 Offsets: Uniform** | **0.926** | **0.6349** | **0.9655** | **0.5046** | **0.4831** | 0.9259 | 0.504 | **0.708** |
| **26 Offsets: Feature importance guided** | 0.9246 | 0.6206 | 0.9568 | 0.4425 | 0.4772 | 0.8873 | 0.474 | 0.698 |
| **8 Offsets: Uniform** | 0.9182 | 0.612 | 0.9542 | 0.4147 | 0.4567 | **0.9275** | **0.533** | 0.652 |
| **8 Offsets: Feature importance guided** | 0.925 | 0.6202 | 0.9564 | 0.4446 | 0.479 | 0.8796 | 0.466 | 0.705 |

**Figure 5.**
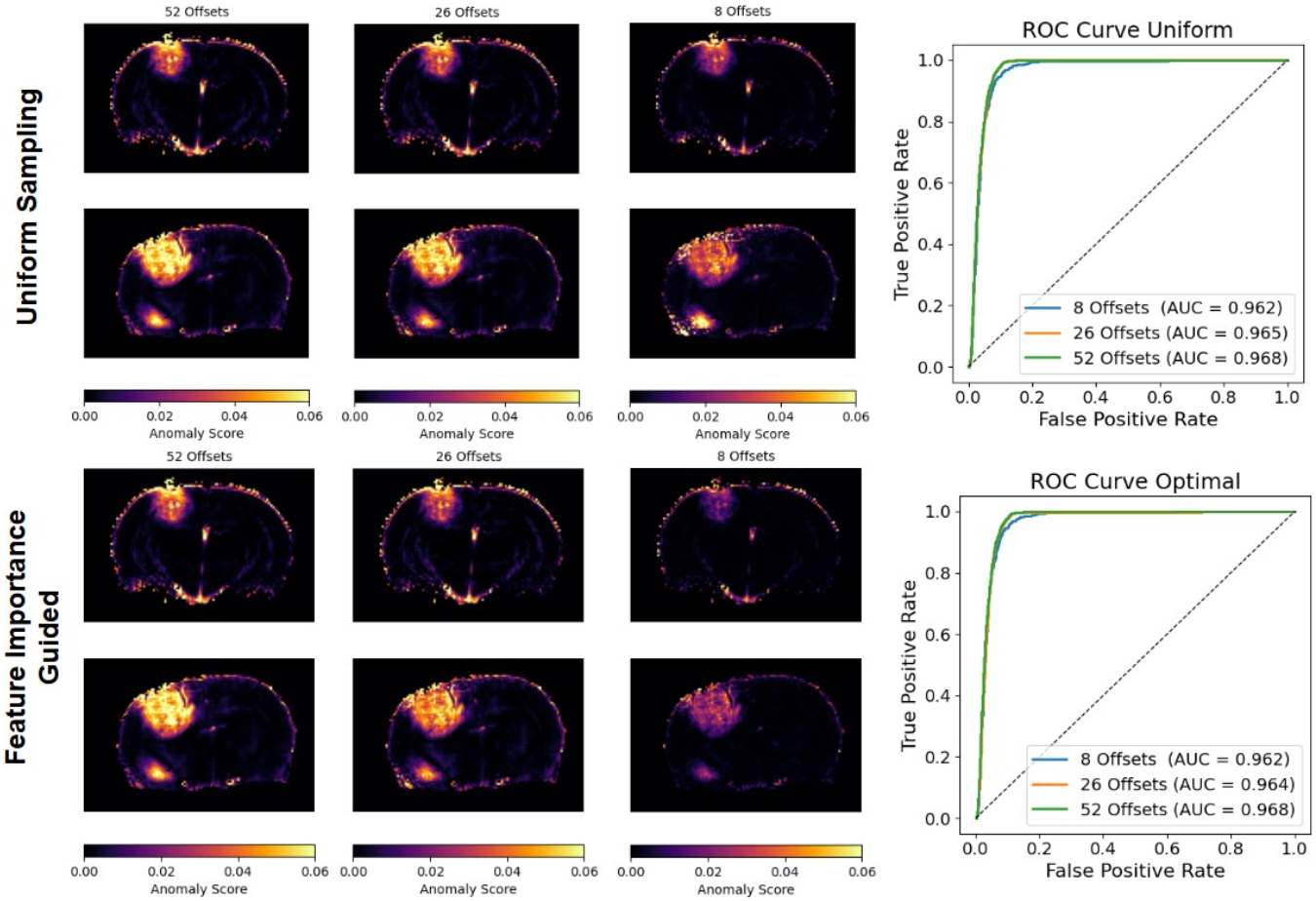
The figure above illustrates the anomaly score maps, and their respective ROCs generated by training the CAE with uniform (a) and feature importance guided (b) sampling schemes, respectively. The figure compares the robustness of the model when sparsely sampled at 52, 26 and 8 offsets, respectively, reaching acceleration factors of 2x and ∼7x.

Furthermore, the feature importance guided scheme seems to overestimate the tumor boundary, likely contributing to higher false positive voxels. This trend is reversed with the 7x acceleration factor, with the uniform scheme overestimating the tumor region. Overall, the results indicate that the quality of anomaly detection is preserved when features are sparse, with optimal sub sampling schemes facilitating the use of anomaly detection in clinical spaces and settings with scan-time constraints.

## Discussion

In this paper, we introduced an unsupervised, voxel-wise anomaly detection pipeline for delineating glioblastoma pathology in tumor bearing rats using CEST MRI Z-spectra post-Gd-enhancement. Unlike prior unsupervised anomaly detection frameworks that operate on structural brain MRI, our model operates on Z-spectra, enabling sensitivity to metabolic changes that may precede or extend beyond morphological abnormalities. Since the model was trained on healthy voxels, it learned to reconstruct an input that aligned with the distribution of healthy Z-spectra. Any spectral pattern that deviated from the reconstruction, as estimated by the mean square error across offsets, was identified as an anomaly. Our findings demonstrate that this biochemically informed, label-free strategy can successfully delineate tumor regions, with Dice coefficients of ∼0.5 and ROC-AUC values greater than 0.95 even when the Z-spectrum is heavily under sampled. When compared to the baseline anomaly detection models, our methodology performed better in identifying anomalies across the majority of metrics. However, IF, when trained using offsets as features similar to our CAE, performed well in detecting anomalous voxels, with an AUC similar to that of the CAE. However, our CAE offers advantages over IF as it can provide feature level anomaly maps (i.e., an anomaly map per offset). Since each offset encodes different metabolic information (-3.5ppm corresponds to rNOE while +3.5ppm corresponds to APT), identifying offsets with the highest reconstruction error on a voxel-wise basis may provide further information on which metabolite is driving a heterogeneous disease phenotype in a spatially consistent manner. Furthermore, the CAE is more robust to noise and nuisance signals, which the Isolation Forest may flag as anomalous, and captures a richer representation of the Z-spectrum, generating a latent space that can be used in downstream tasks to aid in anomaly detection or supervised classifiers.

A significant advantage of our method over existing anomaly detection schemes is incorporation of biochemical information inherent in the Z-spectrum (34). Following the calculation of Shapley values and integrated gradients, we were able to determine offsets that contributed most to the anomaly detection task. The identified offsets (±3.0 to ±5.0ppm from the CAE; -3.5 and -1.2 ppm from IF) reflect contributions from the MT and rNOE pools, with additional APT signal. Correspondingly, ROC-AUC and PR-AUC scores generated from multi-pool fitting derived amplitudes show high values for MT and rNOE, along with high F1-scores and precision. In contrast, APT shows low values across all metrics (**Supplementary Figure 1 and Supplementary Table 1)**. Several studies have demonstrated the utility of MT in distinguishing tumor regions and boundaries, with MT imaging offering advantages over traditional contrast-enhanced T_1_-weighted anatomical images (35, 36, 37). Interestingly, when comparing the reconstruction of healthy and tumor voxels, the residuals for healthy voxels remain largely flat across offsets, while the error for tumor voxels has a broad, Lorentzian-like line shape. This broad shape further suggests that the MT pool is driving the reconstruction error (i.e., anomaly score) observed in the tumor region. Recent studies have also shown changes in rNOE (38, 39), which is sensitive to lipid and protein content, in glioblastoma. However, the feature attributions of offsets related to APT, which has shown clinical utility in tumor delination (40, 41), along with analysis of multi-pool fitting suggest that protein changes may not be the most discriminative metabolic feature. Consequently, the current results indicate that lipid changes may play a larger role in glioblastoma progression and tumor delineation (42, 43, 44). DS showed high ROC-AUC and PR-AUC, but low F1-score and precision. Changes in DS are expected in tumor regions due to changes in T_1_ and T_2_ of bulk water in tumor cells (45, 46). However, the low F1-score and precision indicates that DS may not be a sufficient discriminatory feature. Interestingly, amine showed metrics comparable to rNOE and MT, with feature attributions highlighting offsets around 2 ppm. Future investigations exploring changes in amine contrast in glioblastoma are warranted, as altered amine contrast can arise from pH, tumor infilitration, and different saturation paridigms (i.e., low saturation power v. high saturation power).

Determining the feature importance allowed us to understand the primary offsets contributing to the anomaly score and guide feature selection when sub sampling the Z-spectrum, thus allowing scan-time reduction of prospective experiments that require careful delineation of tumor pathology. Since the acquisition of a Z-spectrum can be on the order of ∼10 – 15 min, reducing the scan time can also facilitate clinical translation, particularly when scan times are limited and dependent on patient comfort. Results from sub-sampling indicate that the quality of anomaly detection is maintained when offsets (i.e., features) are sparse, with metrics extremely close to those of the fully sampled Z-spectrum with the uniform 2x under sampling performing the best. Compared to feature importance guided selection of offsets, the uniform sampling maintains the overall shape of the Z-spectrum. However, when increasing the under sampling to 7x, the feature guided scheme outperforms the uniform scheme, owing to MT’s large contribution to the anomaly score in tumor voxels. Furthermore, uniformly under sampling at such a high acceleration rate may “miss” important offsets around 0 ppm and negatively impact the anomaly score and tumor delineation.

Another major advantage of our approach is that it supplements MPLF by highlighting voxel level spectral deviations independent of explicit pool parametrizations. Although our method does not provide metabolite concentrations, this did not affect subsequent anomaly detection. Our baseline models using fitted parameters as input performed worse compared to the CAE, suggesting that parameter combinations obtained from fitting do not encode sufficiently discriminative information for anomaly detection. This may also indicate an issue with the fitting procedure, and better fitting methods may need to be explored and implemented. Finally, although not unique to our technique, unsupervised anomaly detection requires only healthy data for training which reduces the need for large, labeled datasets. This proves beneficial in heterogenous pathologies or case studies, such as leukodystrophy, in which ground truth labels are hard to obtain and anomalies are rare. Furthermore, anomaly detection can be implemented on an individual basis, allowing for the monitoring of disease progression and precluding the need for statistical analysis that requires large sample sizes and patient stratification.

Despite the proposed advantages of our method, a few notable limitations warrant consideration. Our initial evaluation was performed in a small cohort of tumor-bearing rats, and future work will require validation in a larger cohort with more disease variation. We also did not have access to immunohistochemical results, which makes it challenging to determine diffuse pathological changes and the exact delineation of tumor core and boundary regions. For future investigations, we plan to explore additional network architectures that have shown success in anomaly detection and compare them to our proposed methodology. Our current CAE presents the primary advantages of low computational complexity, stable training, and local inductive bias. Finally, we plan to apply our methodology to pathologies that present more subtle anomalies, such as multiple sclerosis, and demonstrate its performance on human patients, thus bolstering its potential clinical utility. We will expand on our network design by incorporating spatial information, which will enhance the detection of these local anomalies, and include additional Z-spectral information from multiple saturation powers and durations to broaden the sensitivity of our technique to a wider range of metabolites.

Overall, this study establishes unsupervised anomaly detection in CEST MRI as a promising avenue for delineating tumor pathology. By leveraging the biochemical information in Z-spectra, our methodology offers an interpretable, metabolically informed approach for identifying tumor tissue and detecting intra-tumoral heterogeneity, thereby guiding downstream radiologic or therapeutic assessments.

## Acknowledgements

Research reported in this publication was supported by the National Institute of Biomedical Imaging and Bionegineering of the National Institutes of Health under award number P41 EB029460 and by the National Institute on Aging of the National Institutes of Health under award numbers R01 AG063869, RF1 AG087306, and R01 AG091760.

## Data and code availability statement

The data and code used for the generation of results in this manuscript can be found on https://github.com/Abeermathur7/GlioblastomaRatMultiOffset-UAD.git.

## Notes

https://github.com/Abeermathur7/GlioblastomaRatMultiOffset-UAD.git

### Competing Interest Statement

The authors have declared no competing interest.

