## Supplementary Information for "Unsupervised anomaly detection for tumor delineation in a preclinical model of glioblastoma using CEST MRI"

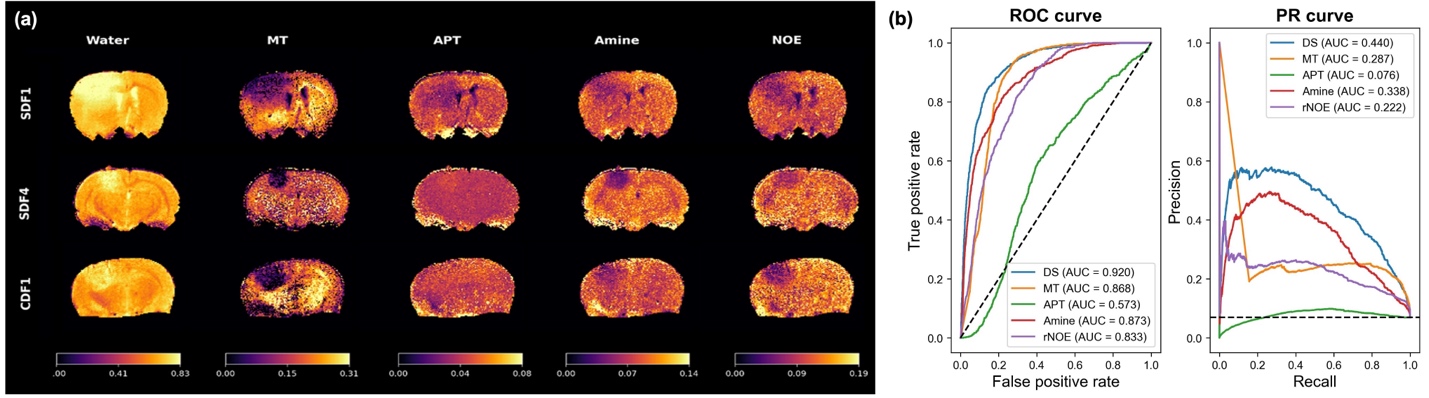

| Metabolite | DS | MT | APT | Amine | rNOE |
| --- | --- | --- | --- | --- | --- |
| F1-score | 0.13 | 0.335 | 0.13 | 0.335 | **0.338** |
| Precision | 0.069 | 0.244 | 0.07 | 0.244 | **0.247** |
| Recall | 0.997 | 0.535 | **1.000** | 0.535 | 0.534 |

**Supplementary Figure and Table 1**: (a) Fitted metabolite maps for five metabolite pools (Direct Saturation, MT, APT, Amine, NOE) on the test set (SDF4 and CDF1) and the validation set (SDF1) along with corresponding ROC curves and AUCs (b). The table below the figures provides the F1-scores for the different metabolites, with rNOE showing the highest F1-score followed by MT and APT.

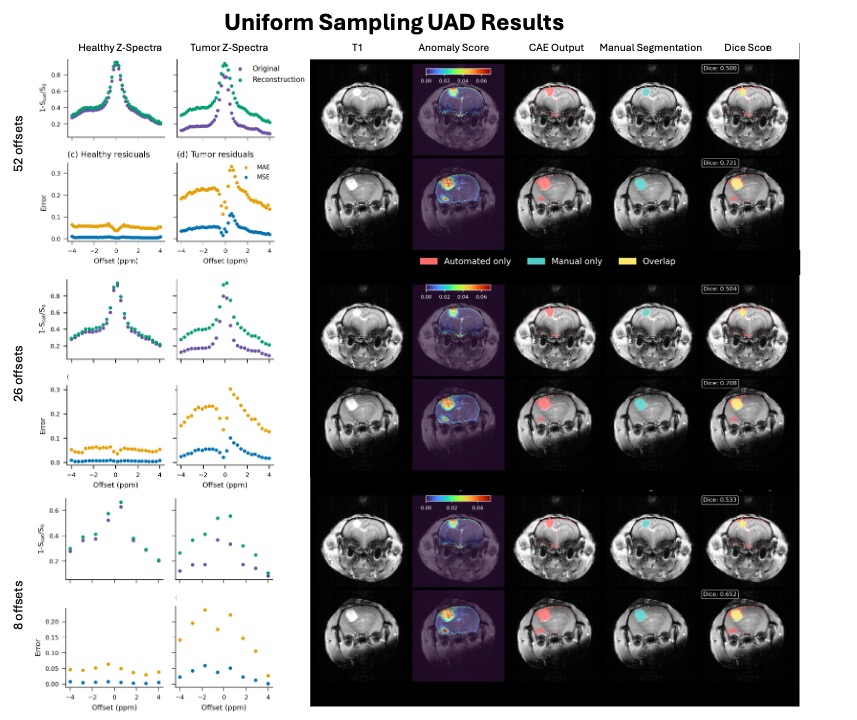

**Supplementary Figure 2:** The figure shows uniformly under sampled data at 2x (26 offsets) and 7x (8 offsets) acceleration factors. The first column depicts the mean Z-spectrum and mean convolutional autoencoder (CAE) reconstruction for healthy and tumor voxels. The second column depicts the reconstruction errors between the original spectrum, with reconstruction errors consistently higher for tumor voxels than healthy and contributing to high anomaly scores as displayed in the anomaly maps. Dice scores indicate high overlap between predicted binary and ground truth tumor masks.

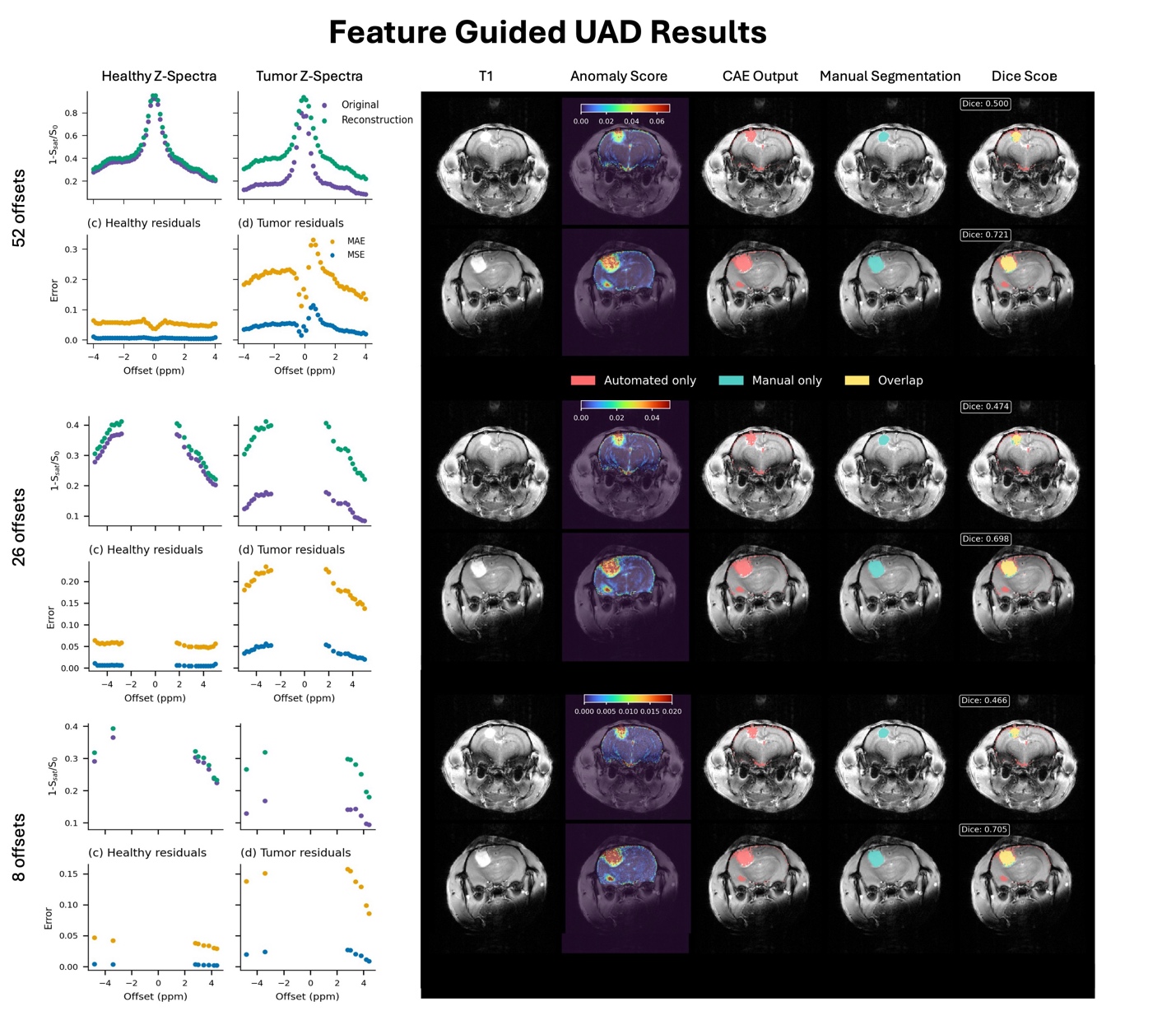

**Supplementary Figure 3:** The figure shows feature guided under sampled data at 2x (26 offsets) and 7x (8 offsets) acceleration factors. The first column depicts the mean Z-spectrum and mean convolutional autoencoder (CAE) reconstruction for healthy and tumor voxels. The second column depicts the reconstruction errors between the original spectrum, with reconstruction errors consistently higher for tumor voxels than healthy and contributing to high anomaly scores as displayed in the anomaly maps. Dice scores indicate high overlap between predicted binary and ground truth tumor masks.
